# A Co-Evolutionary Multi-Objective Framework for Joint Topology and Parameter Discovery in Gene Regulatory Networks

**DOI:** 10.1101/2025.07.22.666158

**Authors:** Tuan Do, Nhung Duong, Luong Doan, Tien Nguyen, Ngoc Do, Anh Truong, Lap Nguyen

## Abstract

Inferring gene regulatory networks (GRN) poses a complex discrete-continuous, multi-objective optimization problem that requires the simultaneous discovery of both network topology and kinetic parameters. Current methods are often limited because they address these two challenges separately. We propose a novel co-evolutionary multi-objective framework that evolves topology and parameter populations in tandem to optimize three competing objectives: dynamic fidelity, network simplicity, and robustness. This algorithm utilizes biologically-inspired operators and is grounded in theoretical results on Pareto optimality for mixed discrete-continuous spaces. In computational experiments on synthetic benchmarks, our approach showed superior performance, attaining the lowest mean squared error in three of four test cases against single-objective and topology-fixed methods. The framework successfully discovers multiple functionally equivalent network configurations, a finding that highlights the degeneracy inherent in biological systems and offers practical insights for synthetic biology, where function is often prioritized over precise structural accuracy.

## 1 Introduction

Gene regulatory networks (GRNs) present a challenging discrete-continuous multi-objective optimization problem where network topology constitutes the discrete combinatorial structure and kinetic parameters form the continuous space [14]. The multi-objective nature arises from balancing data fidelity, network simplicity, and system robustness [12,15].

Traditional approaches assume known topology and focus on single-objective parameter fitting [20,8], yet many applications require simultaneous topology-parameter inference under multiple objectives [6,16]. We propose a co-evolutionary multi-objective algorithm where topology and parameter populations evolve in tandem [2], employing domain-specific operators [11,17] to balance model accuracy, simplicity, and robustness.

### 1.1 Key Contributions

We introduce a discrete-continuous multi-objective framework treating both network structure and parameters as decision variables through a dual-population co-evolutionary approach. The theoretical foundation establishes Pareto-optimal solution existence via Lipschitz continuity and boundedness assumptions [19,1]. Computational experiments on synthetic benchmarks demonstrate superior performance over single-objective and topology-fixed methods, achieving improved accuracy with reduced network complexity.

## 2 Literature Review

### 2.1 Multi-Objective Optimization in Systems Biology

Multi-objective optimization has become critical in systems biology due to inherently conflicting objectives including data fidelity, biological plausibility, and robustness [12,15,16]. Evolutionary algorithms are particularly suited for the large, non-convex, discrete-continuous search spaces characteristic of biological networks [18].

### 2.2 Co-Evolutionary and GRN Inference Methods

Co-evolutionary algorithms partition interdependent populations to optimize coupled components simultaneously [2], naturally capturing biological system interactions. GRN inference typically decomposes into structure discovery and parameter estimation [6,10], with most approaches addressing these separately despite their interdependence [14,24,13].

Current parameter estimation methods assume known topology and employ single-objective optimization [20,8], while structure inference methods often ignore parameter constraints [5,3]. Advanced approaches incorporate biological motifs [11,17] but rarely address joint optimization under multiple objectives with theoretical convergence guarantees [1,19].

## 3 Problem Formulation

We define the decision variables as consisting of two interconnected components: the network topology *E* ⊆ *V* × *V* representing a directed graph among *n* genes, where edges (*j* → *i*) indicate that gene *j* regulates gene *i*, and the parameter set *Θ*(*E*) = {*θ*_*j*→*i*_ : (*j* → ∈ *i*) *E*}, where each *θ*_*j*→*i*_ encompasses biologically relevant parameters such as Hill coefficients and reaction rate constants.

The synthetic dynamics are governed by a continuous-time dynamical system described by the differential equation [10]:

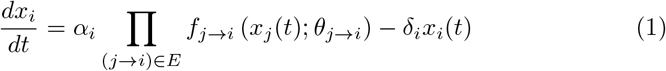

where *x*_*i*_(*t*) represents the concentration or expression level of gene *i* at time

*t*. Each regulatory function *f*_*j*→*i*_ is typically implemented as a Hill-type function bounded within the interval [0, 1] and parameterized by *θ*_*j*→*i*_.

The multi-objective framework addresses three fundamental biological objectives denoted as **F** = (*F*_1_, *F*_2_, *F*_3_). The first objective *F*_1_ quantifies data-fitting error by measuring how well the model matches target synthetic time-series data 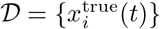. The second objective *F*_2_ addresses network simplicity by penalizing both large numbers of edges |*E*| and extreme parameter values that may be biologically unrealistic. The third objective *F*_3_ measures robustness by quantifying the fraction of initial-condition or parameter perturbations that preserve the characteristic dynamic behavior [16].

The discrete-continuous feasible set is formally defined as:

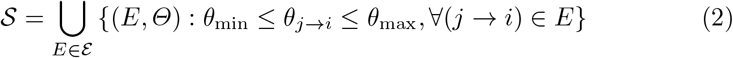

where *ε* represents the space of all feasible topologies, potentially constrained by maximum in-degree requirements or total edge count limitations to maintain biological realism.

A solution (*E*^′^, *Θ*^′^) dominates another solution (*E, Θ*) if *F*_*k*_(*E*^′^, *Θ*^′^) ≤ *F*_*k*_(*E, Θ*) for all objectives *k*, with strict inequality holding for at least one objective [19]. The Pareto set 𝒫 ⊂ 𝒮 consists of all non-dominated solutions, representing the optimal trade-offs between the competing biological objectives.

## 4 Co-Evolutionary Multi-Objective Method

### 4.1 Algorithmic Overview

The co-evolutionary framework employs two distinct yet interdependent populations: a topology population 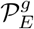 containing candidate regulatory network structures *E*, and a parameter population 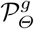 containing parameter sets *Θ* [2]. At each generation *g*, the algorithm proceeds through three main phases.

During the pairing and evaluation phase, each topology *E* from 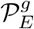 is evaluated in combination with multiple parameter sets *Θ* from 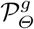. For each topology-parameter pair (*E, Θ*), the objective vector **F**(*E, Θ*) is computed by numerically simulating the GRN dynamics and evaluating the resulting time-series against the target data, complexity metrics, and robustness measures [22].

The selection phase merges all evaluated topology-parameter pairs into a unified pool and ranks them using established multi-objective evolutionary algorithm techniques, such as those employed in NSGA-II [4]. An elite set of non-dominated solutions is retained to seed the subsequent generation, ensuring that promising regions of the solution space are preserved while maintaining diversity across the Pareto front.

The reproduction phase employs specialized operators tailored to the biological domain [11,17]. Topology mutation flips the presence or absence of regulatory edges with probability *p*_mut_ while respecting biological constraints such as prohibitions against self-regulation where appropriate. Parameter crossover operations exchange or average parameter subsets among parent solutions, incorporating domain knowledge about parameter correlations and biological feasibility ranges. Elitism ensures that the best non-dominated solutions discovered across all generations are preserved in the final population.

### 4.2 Theoretical Considerations

Under continuity of objective functions *F*_*k*_ in parameter space *Θ* for fixed topology *E*, and parameter bounds [*θ*_min_, *θ*_max_]^|*E*|^, each objective remains bounded. Since topology space *ε*is finite in practice, the discrete-continuous feasible set 𝒮 admits Pareto-optimal solutions [19]. Standard MOEA convergence results apply under elitism and positive mutation probabilities [1]. Hill-type regulatory functions exhibit Lipschitz continuity on bounded intervals [10], facilitating stability analysis.

### 4.3 Algorithmic Complexity

The combinatorial topology space scales as 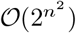 while each topology induces a continuous parameter subspace of dimension 𝒪 (*n*^2^), creating a mixed discrete-continuous optimization challenge. Population-based heuristics are essential as exact enumeration becomes computationally prohibitive for moderate network sizes, requiring efficient exploration of promising regions through multi-objective selection.

## 5 Computational Experiments

### 5.1 Experimental Design

We created benchmark sets containing synthetic GRN instances with known true parameters *Θ*^true^ that generate distinct dynamic behaviors including stable equilibria, oscillatory patterns, and multistable systems. The benchmark collection includes small networks with 5 genes where each gene is regulated by at most 2 others, and medium-sized networks with 10 genes exhibiting moderate connectivity patterns.

Synthetic time-series data 𝒟 is generated by numerically simulating the true GRNs using established ODE integration techniques [22]. The algorithm is then challenged to recapitulate these dynamics through joint topology and parameter search, without prior knowledge of the ground-truth network structure or kinetic parameters.

### 5.2 Algorithm Setup

Population sizes of 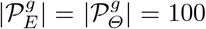 enable adequate exploration of discrete-continuous spaces. Topology mutations occur with probability *p*_mut_ = 0.1 while parameter mutations use probability 0.05 with Gaussian perturbations. Evaluation involves numerical ODE integration over *t* ∈ [0, *T*] with MSE fidelity measurement and systematic robustness analysis [22].

### 5.3 Comparative Methods

We compare against three baselines on 5-and 10-gene networks over 50 generations with 10 independent runs: topology-fixed parameter estimation using ground-truth structure [20,8], random topology search with single-objective fitting, and pure single-objective MSE minimization ignoring complexity and robustness.

### 5.4 Pareto Front Analysis

We implement comprehensive Pareto front analysis examining solution diversity and functional equivalence characteristics [7,9]. A representative 10-gene instance (benchmark size10 idx2) provides detailed case study analysis over 50 generations. Key solutions include: best dynamic fidelity (minimal MSE), highest robustness (maximal perturbation resistance), and simplest architecture (minimal complexity penalty). Analysis quantifies regulatory degeneracy where distinct topologies produce identical dynamics [21], examines motif conservation patterns [11], and tracks convergence dynamics to validate population diversity maintenance throughout optimization.

## 6 Results and Analysis

### 6.1 Comparative Performance

Table 6.1 presents a comprehensive performance comparison across methods and benchmarks, revealing several key trends in the relative performance of different approaches to GRN inference.

### 6.2 Case Study: Pareto Front Analysis

We analyze a representative 10-gene optimization (benchmark size10 idx2) demonstrating multi-objective trade-offs and functional equivalence discovery. The optimization convergence analysis reveals several important characteristics of the co-evolutionary search process across four key metrics [4,2]. The best data-fitting error (MSE) shows rapid initial improvement followed by more gradual refinement, achieving values below 0.01 by generation 20 and continuing to improve throughout the optimization process. Network complexity, measured by average edge count, exhibits controlled fluctuation around 13-14 edges, demonstrating the algorithm’s ability to balance network parsimony with functional requirements. The robustness metric remains consistently high (approximate 1) throughout optimization, indicating that discovered solutions maintain stable behavioral characteristics even as the algorithm explores different topological configurations. Solution diversity, tracked by Pareto front size, varies between 20-38 solutions throughout optimization, demonstrating sustained population diversity crucial for revealing functionally equivalent networks.

**Table 1.**
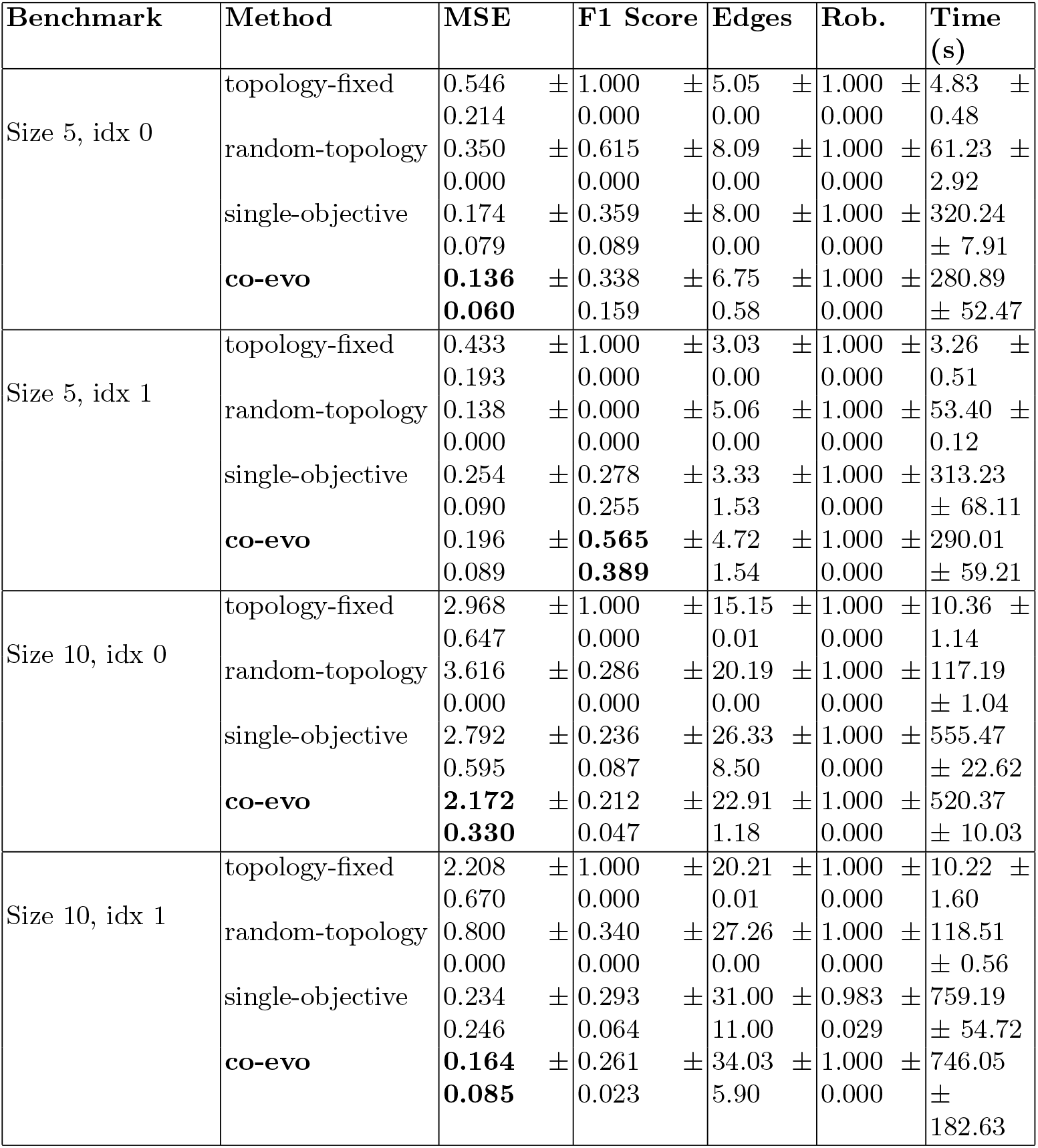
Performance Comparison Across Methods and Benchmarks.

The best MSE solution (Figure 2) achieves remarkable fidelity (MSE = 0.001123) using only 23 edges versus 27 in the ground truth. Gene expression dynamics comparison reveals near-perfect reproduction of target behaviors across all 10 genes [10,22]. The predicted trajectories closely match target dynamics for both monotonic decay patterns (Genes 0-4, 6-8), transient responses, and non-monotonic patterns (Gene 5), with the co-evolutionary approach successfully discovering functionally equivalent regulatory architectures that accurately reproduce complex dynamic behaviors including oscillatory and steady-state responses. The F1 score of 0.320 indicates moderate structural accuracy, yet functional performance suggests a viable alternative regulatory architecture. Parameter values show appropriate diversity with Hill coefficients ranging from 1.2 to 4.2 and dissociation constants spanning 1.5 to 9.3, reflecting biological heterogeneity. This demonstrates functional equivalence where distinct regulatory architectures achieve identical dynamic behaviors through different molecular mechanisms [7,9].

**Fig. 1.**
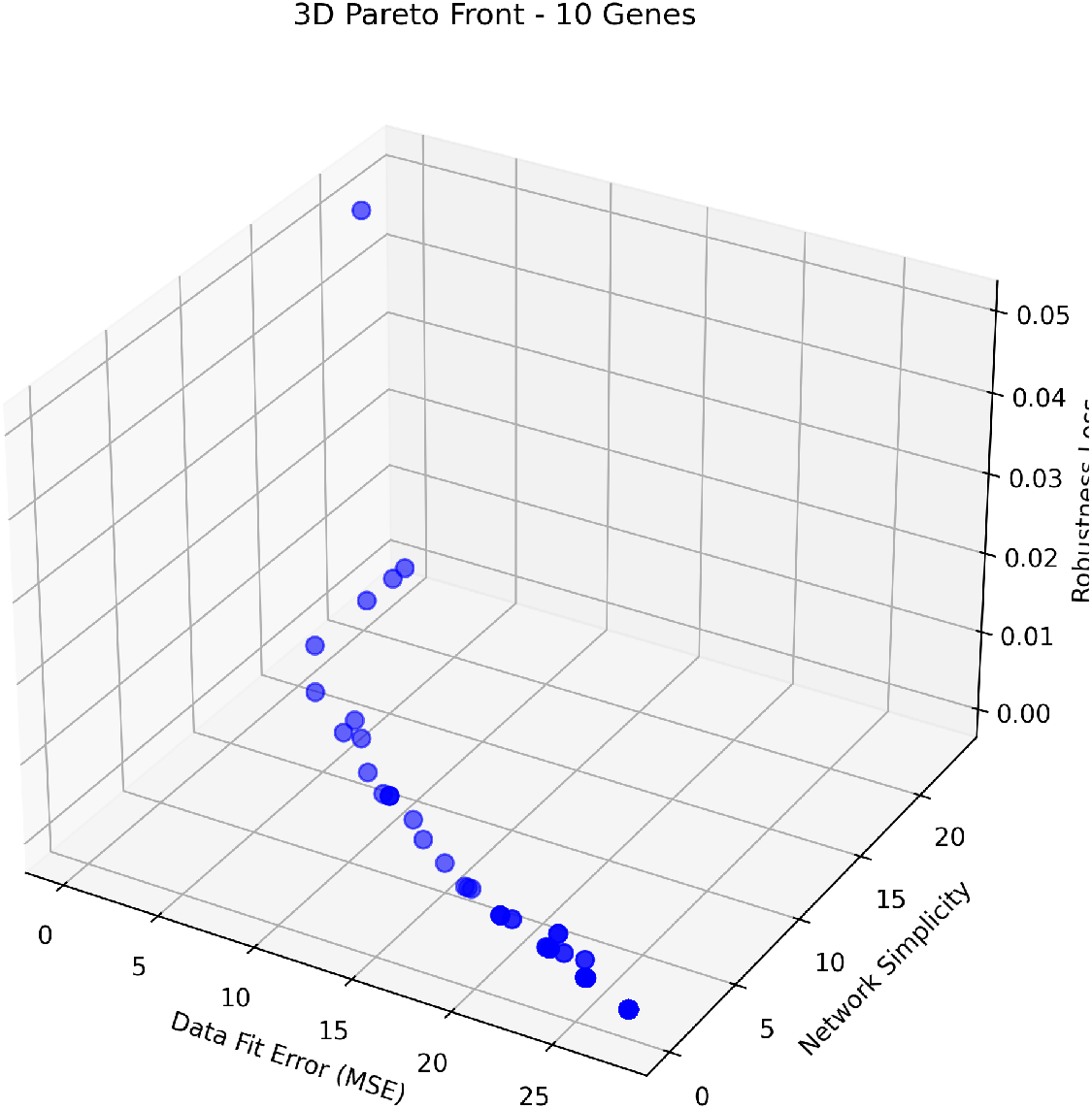
Three-dimensional Pareto front showing 25 non-dominated solutions across data-fitting error, network simplicity, and robustness objectives.

**Fig. 2.**
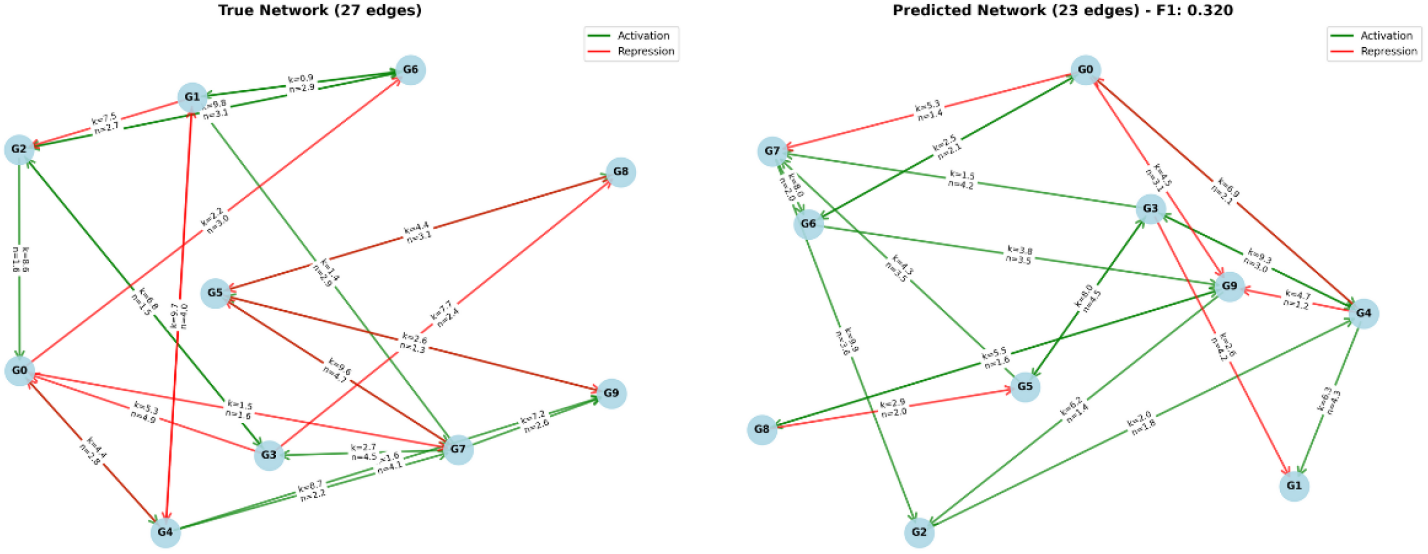
Network topology comparison: (left) ground-truth network with 27 edges and (right) predicted network with 23 edges achieving F1 score of 0.320 and MSE of 0.001123.

## 7 Discussion

### 7.1 Implications for Systems Biology

Our results demonstrate that joint topology-parameter inference using co-evolutionary multi-objective optimization discovers networks reproducing target dynamics more accurately than methods addressing components separately [14,6]. Lower F1 scores relative to topology-fixed methods reflect fundamental biological realities: systems frequently exhibit structural degeneracy where multiple topologies produce identical dynamics [7,9,21,23]. This functional equivalence represents a key characteristic of biological systems achieving outcomes through different molecular mechanisms.

The multi-objective framework naturally prioritizes functional equivalence over exact structural matching when alternative topologies offer superior overall performance [12,15,16]. This reflects realistic biological design priorities where structural efficiency and robustness often dominate specific interaction patterns. Limited time-series data creates identifiability challenges where multiple configurations explain dynamics equally well [24,13,5,3]. Our approach navigates this uncertainty by identifying robust, parsimonious solutions satisfying different biological design criteria.

For synthetic biology, the framework reveals multiple viable designs achieving identical goals while varying in complexity and implementability [11,17], providing engineers flexibility based on practical constraints such as available genetic components or metabolic burden.

### 7.2 Methodological Advantages

The co-evolutionary structure enables balanced exploration of diverse networks while refining parameters within promising topological regions [2]. This dual-population approach prevents premature convergence in the combined discrete-continuous space. Theoretical foundations rest on established multi-objective optimization results [1,19], with Pareto-optimal solution existence guaranteed under continuity and boundedness assumptions. Lipschitz continuity of Hill-type functions [10] provides stability guarantees.

Despite combinatorial complexity scaling as 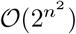, the approach efficiently navigates search space through multi-objective evaluation. Domain-specific operators [11,17] enhance biological plausibility while maintaining mathematical rigor. The multi-objective formulation captures inherent trade-offs more accurately than single-objective approaches focusing solely on data-fitting [20,8].

### 7.3 Limitations and Future Work

Computational demands limit application to larger networks, requiring parallel implementation or surrogate modeling approaches. Current robustness metrics capture limited aspects of biological stability [15]; future work should investigate alternative formulations including dynamic stability analysis and multiperturbation sensitivity. Real biological data introduces challenges including noise, partial observations, and experimental variability [22], requiring noise-robust objectives and uncertainty quantification.

The framework should be extended to complex dynamics including multi-stable systems and chaotic behavior [3], and integrated with experimental design methodologies to reduce identifiability challenges through strategic data collection.

## 8 Conclusion

This paper introduces a novel co-evolutionary multi-objective framework for joint topology and parameter discovery in synthetic gene regulatory networks. Through simultaneous evolution of regulatory network structures and their corresponding kinetic parameters while balancing multiple biologically relevant objectives, our approach discovers networks that more accurately reproduce target dynamics compared to baseline methods, while maintaining reasonable complexity and robust behavioral characteristics.

Our results demonstrate that the pursuit of perfect topological matching (high F1 scores) may be less critical than identifying functionally equivalent networks that achieve desired behaviors with appropriate complexity and robustness characteristics. This perspective aligns with practical needs in systems biology and synthetic biology, where functional equivalence often carries greater importance than structural identity.

The co-evolutionary multi-objective approach provides a valuable tool for realistic GRN discovery scenarios where both topology and parameters remain unknown, and multiple competing objectives must be balanced simultaneously. By integrating discrete topology search with continuous parameter optimization within a theoretically grounded framework, our method addresses an important gap in the systems biology toolkit and demonstrates the value of operations research techniques in biological system design.

The framework reveals multiple functionally equivalent network configurations, reflecting a phenomenon observed in natural systems as developmental canalization [21,23]. By prioritizing functional equivalence over exact structural reconstruction, our approach provides valuable insights for synthetic biology design where implementing precise regulatory behavior often outweighs the need for specific molecular interactions. Additionally, the method identifies critical regulatory connections that remain essential across multiple solution candidates, suggesting evolutionarily conserved interaction patterns.

Future work will focus on extending the approach to larger networks, incorporating more sophisticated robustness measures, and adapting the framework for real biological data with associated experimental uncertainties. The demonstrated success on synthetic benchmarks provides a strong foundation for these extensions and suggests broad applicability for biological network inference and design challenges.

## References

1. Abraham, A., Jain, L.: Evolutionary multiobjective optimization. Springer (2005)

2. Antonio, L.M., Coello, C.C.C.: Coevolutionary multiobjective evolutionary algorithms: Survey of the state-of-the-art. IEEE Transactions on Evolutionary Computation 22 (2018) 851–865

3. Daneker, M., Zhang, Z., Karniadakis, G., Lu, L.: Systems biology: Identifiability analysis and parameter identification via systems-biology informed neural networks. Methods in molecular biology 2634 (2022) 87–105

4. Deb, K., Agrawal, S., Pratap, A., Meyarivan, T.: A fast and elitist multiobjective genetic algorithm: Nsga-ii. IEEE Trans. Evol. Comput. 6 (2002) 182–197

5. Degasperi, A., Fey, D., Kholodenko, B.: Performance of objective functions and optimisation procedures for parameter estimation in system biology models. NPJ Systems Biology and Applications 3 (2017)

6. den Broeck, L.V., Gordon, M., Inzé, D., Williams, C.M., Sozzani, R.: Gene regulatory network inference: Connecting plant biology and mathematical modeling. Frontiers in Genetics 11 (2020)

7. Green, S., Batterman, R.W.: Making sense of top-down causation: Universality and functional equivalence in physics and biology. In: Top-down causation and emergence. Springer (2021) 39–63

8. Gabor, A., Banga, J.: Robust and efficient parameter estimation in dynamic models of biological systems. BMC Systems Biology 9 (2015)

9. Klein, B., Swain, A., Byrum, T., Scarpino, S., Fagan, W.: Exploring noise, degeneracy and determinism in biological networks with the einet package. Methods in Ecology and Evolution 13 (2022) 799 – 804

10. Likhoshvai, V.A., Ratushny, A.: Generalized hill function method for modeling molecular processes. Journal of bioinformatics and computational biology 5 2B (2007) 521–31

11. Liu, W., Sun, X., Yang, L., Li, K., Yang, Y., Fu, X.: Nscgrn: a network structure control method for gene regulatory network inference. Briefings in bioinformatics (2022)

12. Malashin, I.P., Martysyuk, D., Tynchenko, V., Gantimurov, A.P., Nelyub, V., Borodulin, A.S.: Integrating machine learning and multi-objective optimization in biofuel systems: A review. IEEE Access 13 (2025) 81983–82002

13. Meshkat, N., zhen Kuo, C.E., DiStefano, J.: On finding and using identifiable parameter combinations in nonlinear dynamic systems biology models and combos: A novel web implementation. PLoS ONE 9 (2014)

14. Meyer, P., Cokelaer, T., Chandran, D., Kim, K., Loh, P.R., Tucker, G., Lipson, M., Berger, B., Kreutz, C., Raue, A., Steiert, B., Timmer, J., Bilal, E., Sauro, H., Stolovitzky, G., Sáez-Rodríguez, J.: Network topology and parameter estimation: from experimental design methods to gene regulatory network kinetics using a community based approach. BMC Systems Biology 8 (2014) 13 – 13

15. Nijhout, H., Best, J., Reed, M.: Systems biology of robustness and homeostatic mechanisms. Wiley Interdisciplinary Reviews: Systems Biology and Medicine 11 (2018)

16. Noman, N., Monjo, T., Moscato, P., Iba, H.: Evolving robust gene regulatory networks. PLoS ONE 10 (2015)

17. ou Zhao, Y., Chen, Y., Jiang, M.: Automatic inference of gene regulatory network using dynamic model based on law of mass action. The Journal of Information and Computational Science 12 (2015) 893–904

18. Rawlins, G.J.: Foundations of Genetic Algorithms 1991 (FOGA 1). Volume 1. Elsevier (2014)

19. Roy, A., So, G., Ma, Y.: Optimization on pareto sets: On a theory of multi-objective optimization. ArXiv abs/2308.02145 (2023)

20. Schälte, Y., Fröhlich, F., Jost, P.J., Vanhoefer, J., Pathirana, D., Stapor, P., Lakrisenko, P., Wang, D., Raimúndez-Álvarez, E., Merkt, S., Schmiester, L., Städter, P., Grein, S., Dudkin, E., Dorešić, D., Weindl, D., Hasenauer, J.: pypesto: a modular and scalable tool for parameter estimation for dynamic models. Bioinformatics 39 (2023)

21. Siegal, M., Bergman, A.: Waddington’s canalization revisited: Developmental stability and evolution. Proceedings of the National Academy of Sciences of the United States of America 99 (2002) 10528 – 10532

22. Sommer, A.: Numerical methods for parameter estimation in dynamical systems with noise with applications in systems biology. PhD thesis, Heidelberg University (2017)

23. Spirov, A.V., Sabirov, M.A., Holloway, D.M.: Systems evolutionary biology of waddington’s. Evolutionary Physiology and Biochemistry: Advances and Perspectives (2018) 167

24. Valderrama-Bahamondez, G.I., Fröhlich, H.: Mcmc techniques for parameter estimation of ode based models in systems biology. Frontiers Appl. Math. Stat. 5 (2019) 55

